# Visualization and Prediction of *in vivo* Phosphate Dynamics via Auto-Glowing Plant Sensors

**DOI:** 10.1101/2025.07.30.667770

**Authors:** Ching-Wen Chiu, Ya-Fen Chan, Ting-Yang Lu, Liang-Jie Chiu, Ya-Ru Li, Hao-Jun Lu, Huai-Ji Huang, Alexander S. Mishin, Karen S. Sarkisyan, Chiu-Han Hsiao, Ming-Jung Liu

## Abstract

Monitoring endogenous nutrient levels is crucial for maximizing crop yields and optimizing fertilizer use. Here, focusing on phosphorus, an essential nutrient for plant growth, we developed a low-cost and non-invasive biosensor to visualize and predict early stress signaling in plants. By combining plant phosphate (Pi)-deficiency-induced promoter systems with fungal self-sustained bioluminescence systems genetically engineered into tobacco plants, we created sensor plants that emitted more light when experiencing Pi deficiency. This light emission correlated with the expressions of known phosphate-responsive genes and the total phosphorus content in plants, and decreased during Pi recovery conditions, demonstrating the responsiveness and robustness of the sensor plants in reflecting endogenous phosphorus deficiency. The sensor plants responded primarily to Pi deficiency rather than nitrogen or potassium deficiencies and were sensitive to different ranges of external Pi concentrations. Additionally, when grafted onto tomato and chili pepper plants, the sensor plants responded to external phosphorus deficiency, showing promise for monitoring stress signals in different crop species. Using deep-learning-based image analysis techniques, auto-luminescent signals of sensor plants could be detected and used to predict phosphorus deficiency. This study outlines a strategy of creating a self-luminous biosensor to visualize phosphate dynamics *in planta* and predict nutrient deficiency for sustainable agriculture.

## Introduction

### Detection of nutrient-deficiency stress in plants

Micronutrients such as nitrogen, phosphorus, and potassium are essential for plant growth development and reproduction (Cordell & White, 2014; Sadoine *et al.*, 2023). For example, phosphorus, one of the top nutrient elements, promotes root growth and fruit development, thus affecting crop yields (Holford, 1997). However, climate instability has widely disturbed ecosystems, impacting the soil nutrient imbalance and affecting nutrient availability. Extreme flooding and droughts lead to increased nutrient leaching, runoff, and the change of soil pH values, which can negatively impact soil health and as a result of limiting the nutrient uptake by plants. Additionally, plants obtain phosphorus from the soil mainly in the forms of inorganic phosphate (Pi), by phosphate transporters into the root cells. While the total phosphorus content in soil can be relatively high, due to the low solubility of Pi in soil, the uptake of Pi by plants is often limited (Raghothama, 1999; Paz-Ares *et al.*, 2022). As a consequence, phosphorus limitation is a common condition in many agricultural lands.

To cope with nutrient limitation, nutrient fertilizers are widely applied in agriculture However, only a limited amount of the fertilizer supply is taken up by plants, and the rest is leached into the environment with detrimental effects, such as carbon emission, soil degradation and water quality in lands (Cordell & White, 2014). Thus, the proper usage of nutrient fertilizers is critical for both plant yields and soil health, supporting agricultural production and agricultural sustainability. Various tools such as selective electrodes, optical sensors, reflectometers, and image-based sensors have been developed to quantify macronutrients in plant fields (Raul *et al.*, 2016; Sadoine *et al.*, 2023). Mechanical sensor based on optical and electrochemical techniques is an effective approach to examining soil nutrient concentrations (Burton *et al.*, 2020). Nevertheless, the dynamic changes of external environments may not be directly associated with plant physiological status. Thus, to observe the stress status within living plants directly, reporter genes like β-glucuronidase (GUS), and green fluorescent protein (GFP) have been introduced into plants, allowing monitoring of *in vivo* nutritional status, including magnesium and phosphate levels (Bustos *et al.*, 2010; Kamiya *et al.*, 2012; Lin *et al.*, 2016). Additionally, the advance of hyperspectral imaging techniques and portable spectrometer development enables to detect the physiological and biochemical changes inside plants, inferring abiotic stresses *in vivo* (Burton *et al.*, 2020; Saric *et al.*, 2022; Sadoine *et al.*, 2023). While these reporter genes and imaging/spectra analyses serve as powerful tools for assessing nutritional status, challenges of detecting stress signals and high cost instruments hinder their application in detecting whole plants in field studies (de Ruijter *et al.*, 2003). These include the signal quantification *in vitro* (i.e., the invasive detection), high background and low dynamic signal ranges, the early detection of plant stress signals before severe physiological damages occur and the need for laser excitation and hyperspectral cameras requiring expensive instruments or exogenous chemical addition into plants (de Ruijter *et al.*, 2003). In addition, the spectrum- and imaging-based analyses provide information of general physiological and physical changes (such as metabolites and leaf color) but not specify which abiotic stress that plants are facing (Saric *et al.*, 2022).

Luminescence, a common phenomenon producing light, has been found in living organisms from bacteria to fungi and animals (Fleiss & Sarkisyan, 2019). These bioluminescent reactions depend on the luciferase-luciferin pair, in which luciferase enzyme oxidizes a specific substrate, luciferin, to create light. While the ability of emitting bioluminescence in darkness has been observed among many species, the bioluminescent system has not been reported in terrestrial plants (Fleiss & Sarkisyan, 2019; Li *et al.*, 2021). The feature of bioluminescence with green spectrum and no excitation required provides the advantage of minimizing chlorophyll absorption and a low-noise background in plants (Reuter *et al.*, 2020; Li *et al.*, 2021). The luciferase gene has been initially proven to be a powerful reporter in both transient expression assay and in transgenic plant systems. Upon the external luciferin addition, luciferase-encoding plants produce clear luminescent signals. However, challenges like luciferin substrate costs and penetration into whole plants and tissues have limited its application. Importantly, based on the newly discovered bioluminescent systems and enhancement of suboptimized luminescence, the glowing plants with robust auto-luminescent signals were created (Khakhar *et al.*, 2020; Mitiouchkina *et al.*, 2020; Shakhova *et al.*, 2024). By genetically engineering fungal genes for luciferin synthesis and recycling, along with the fungal luciferase gene, into plant genomes, these plants emit visible autonomous bioluminescence detectable by the naked eye and captured by consumer-grade cameras. (Khakhar *et al.*, 2020; Mitiouchkina *et al.*, 2020). These reports showed the successful reconstitution of a fungal bioluminescence pathway, involving the provision of both luciferin and luciferase, within plants. This integration allows plants to autonomously emit light in darkness. The demonstrated auto-glowing capability suggests that these plants can serve as a natural source of illumination, enabling the reading and writing of signals from plants that can be detected by a low-cost device. The features of self-sustained bioluminescence and high sensitivity make this advancement exciting to develop a plant biological sensor with noninvasive inspection. This type of sensor, coupled with low-cost camera devices, could monitor plant stresses, providing real-time readouts on the physiological status of living plants. This autonomous signal emission capability holds significant potential for practical applications in field studies.

Thus, in this study, we aim to develop a sensor plant with stress-induced auto-luminescent activity, coupled with a machine-learning-based technique, that provides a non-invasive and low-cost tool to monitor endogenous stress status and immediately report stress signals. We focus on phosphorus, one of the most essential macronutrients and in the format of phosphate (Pi) accessible for plants, and chose the promoter of a phosphate starvation-induced gene (*TPSI1*) that has been experimentally validated for driving Pi deficiency-response gene expressions (Lin *et al*., 2016). Tobacco serves as our plant model system, selected for its genetic engineering efficiency, tissue culture adaptability, and grafting compatibility across species (Notaguchi *et al*., 2020). We genetically introduced the fungal bioluminescent systems, coupled with the luciferase gene driven by a Pi-deficiency-induced *TPSI1* promoter, into tobaccos to generate Pi-sensitive auto-luminescent sensor plants. By monitoring plant-emitted auto-luminescence signals via a low-cost camera, we assessed the responsiveness, sensitivity and nutrient-specificity of the Pi-sensitive sensor plants. We also assessed the impact of the bioluminescent system in plant Pi deficiency response of sensor plants. Furthermore, we establish a digital pipeline integrating imaging analysis and deep-learning techniques to predict stress signals. Lastly, to explore the extent that sensor plants can be applied to monitor stress signals on different crop species, we assessed the Pi-deficiency response of the sensor plants when grafting onto target crop species.

## MATERIALS and METHODS

### Constructs for constituting fungal bioluminescence pathways into tobacco

The bioluminescent plasmid generated previously (Mitiouchkina *et al*., 2020) was applied here, which contains five fungal genes from *Neonothopanus nambi*: *NpgA* (phosphopantetheinyl transferase), *HispS* (hispidin synthase), *H3H* (hispidin-3-hydroxylase), *CPH* (caffeoyl pyruvate hydrolase) and *Luz* (luciferase) (**Fig. 1A**). *NpgA, HispS, H3H* and *CPH* genes for luciferin synthesis and recycle were driven by the promoters of either cauliflower mosaic virus 35S, *Arabidopsis thaliana* actin2 or figwort mosaic virus as described previously (Mitiouchkina *et al*., 2020). The 35S promoter sequences from pX037 (Mitiouchkina *et al*., 2020) and *TPSI1* sequences described in a previous study (Lin *et al*., 2016) were cloned into pENTR ™ 5’-TOPO ® vector by TA cloning kit (Invitrogen™, K59120) to generate the 35S- and TPSI1-promoter entry clone. The promoter region of *Luz* gene in the bioluminescent plasmid (mentioned above) was replaced with 35S and *TPSI1* promoter fragments in the 35S and TPSI1 entry clones via LR recombination reaction (Invitrogen™, 12538120). To generate the Vector plasmid (i.e., without promoter sequence), 35S promoter entry clone was digested by *Eco*RI and self-ligated to create an entry clone. The “Vector” entry clone with no promoter region was then cloned into the bioluminescent plasmid via LR recombination reaction as described above.

**Figure 1.**
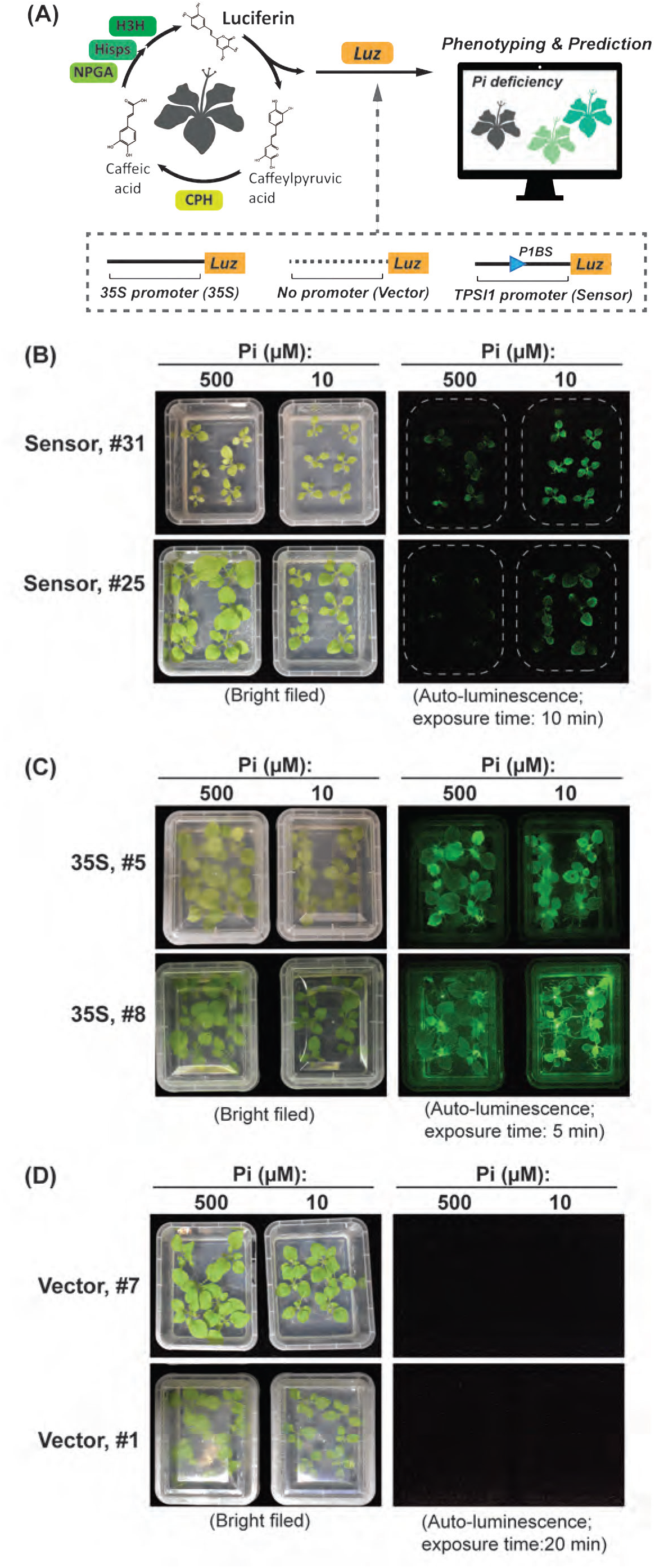
The generation of plant sensors with Pi-deficiency-induced bioluminescent systems. (A) Schematic of generating Pi deficiency-induced auto-glowing sensor plants by integrating fungal bioluminescence pathways and phosphate (Pi) responsive transcriptional regulation. The genes involved in luciferin synthesis and recycling (green boxes) and encoding luciferase (Luz; orange box) were integrated into tobacco genomes. The *Luz* gene is driven by 35S (35S) and Pi-deficiency induced *TPSI1* (Sensor) promoters and without promoter (Vector), respectively. (B) The auto-luminescent signals from the images captured by Nikon D5600 DSLR digital camera in two independent transgenic tobacco lines of 35S, Vector, and Sensor plants. Plants were grown in agar boxes (with full medium) for ∼ 10 days and subjected to Pi sufficiency (500 uM Pi) and Pi deficiency (10 uM) treatments Pi, respectively, for another 10 days.

### Plant growth, grafting and nutrient-deficiency treatment

The tobacco (*Nicotiana benthamiana*) plants are grown on vermiculite and soil in a growth chamber under a 14-h light (8:00–22:00, 150 μmol m^−2^s^−1^)/10-h dark cycle at 25°C and cultured with half-strength modified nutrient solution (Hoagland & Arnon, 1950; Millner & Kitt, 1992) for 18∼20 days before the specified nutrient treatments. The soils with limited phosphorus were excavated from the deep layers of the field (**Supplement Table S1**). For the phosphate deficiency/sufficiency/recovery experiments, the tobacco seedlings were supplemented with the half-strength modified nutrient solution with the adjustment of KH_2_PO_4_ concentrations: 500 µM (+Pi, Pi-sufficient/-recovery) and 0 µM(−Pi, Pi-deficient) as described previously (Lin *et al*., 2016). For the test of the Pi-deficiency sensitivity, the tobacco seedlings were supplemented with nutrient solutions with 500 µM, 200µM, 50µM, 10µM, and 0 µM KH_2_PO_4_. The duration of nutrient treatments was indicated within the given figures and not mentioned here.

For nutrient-specific deficiency experiments, the tobacco seedlings were maintained in a half-strength modified nutrient solution with the original full nutrients (Hoagland & Arnon, 1950; Millner & Kitt, 1992) and transferred to a solution in which specific nutrient elements were omitted. To compensate for the ion concentrations, an equivalent concentration of ions was supplied. The details of nutrient-specific deficiency treatment were as follows: for -N (nitrate-deficiency), Ca(NO_3_)_2_ and KNO_3_ were replaced with CaCl_2_ and KCl; for -P (phosphate-deficiency) treatment, KH_2_PO_4_ was replaced with KCl; for -K (potassium-deficiency) treatment, KNO_3_ and KH_2_PO_4_ were replaced with Ca(NO_3_)_2_ and NaH_2_PO_4_ (Lin *et al*., 2016).

For grafting the tobacco sensor plants onto the target crops, ∼ 30-day-old tobacco plants serve as scions whereas the 30-35-day-old tomato (*Solanum lycopersicum*, cultivar Micro-Tom) and chili pepper (SV-015 Hot pepper) serves as the rootstock. After the cleft grafting for 11-13 days, the plants were supplemented with 500 µM and 0 µM KH_2_PO_4_ for the phosphorus deficiency treatments.

### Screening of transgenic tobacco plants with auto-luminescent system

The transformation of plasmids into tobacco via *Agrobacterium*-infiltration technique was adopted from a previous study (Dandekar & Fisk, 2005). The T0/T1 seed was grown in the resistant medium plates (50 ppm Kanamycin + half-strength modified nutrient solution) for ∼10 days to select the transgenic lines with >75% resistant rate. The selected lines were cultured in the tissue box with the half-strength modified nutrient solution with the concentrations of 500 µM (+Pi, Pi-sufficient/-recovery) and 10 µM (−Pi, Pi-deficient) for ∼14 days for the detection of Pi-deficiency induced auto-luminescent signal by the digital camera. In addition, the selected lines were also subjected to genomic DNA purification to confirm the insertion of bioluminescence-related genes. For genomic DNA purification, the young-leaf tissue of the target lines in the tube was added with 0.5N NaOH buffer and grounded thoroughly at room temperature. The supernatant was transferred to a tube containing 50 μL of 50 µM Tris pH 8.0, mixed thoroughly and used as the PCR templates. The PCR amplification assay was performed in 20μL reaction containing PCR templates, 1x PCR Buffer (Ampliqon taq 2x master mix), 10 μM of forward/reverse primers listed in **Supplemental Table S2**.

The Nikon D5600 DSLR camera with a NIKKOR Z 24-70mm F4S lens was used to capture all photos presented here. Depending on the experimental setup, lens aperture and other considerations, a range of ISO values from 200 to 1600 was used, with exposure times from 1/50 s to 900 s. The photos in dark were captured with an exposure time of 600 seconds and ISO 1600.

### Phosphate (Pi) content assay

The cellular Pi content was determined as described previously (Ames, 1966; Chiou *et al*., 2006). Briefly, fresh leaves (10-50 mg) collected from the Pi-deficiency and sufficiency-treated plants were homogenized and mixed with 0.5 mL of 1% glacial acetic acid, followed by an incubation at 42°C for 30 minutes. The solution underwent centrifugation at 13,000g for 5 minutes, and the supernatant aliquot was utilized in the Pi assay. For the Pi assay, 10 μl of the sample was mixed with 140 μl of the assay solution (0.3% (NH_4_)_6_Mo_7_O_24_ · 4H_2_O, 0.86 N H_2_SO_4_, and 1.4% ascorbic acid) and 50 μl of 1% glacial acetic acid, followed by an incubation at 42°C for 30 minutes. The Pi content was measured at A820 via BioTek CYTATION 5.

### Analysis of mRNA expressions and Gene Ontology (GO) enrichment

Total RNA purification from the fresh leaves of the Pi-deficient-treated plants, the cDNA syntheses, and the quantitative RT-PCR assays were described previously (Chiu *et al*., 2022). The mRNA abundances of specific target genes were determined using the ΔCt (threshold cycle) method, with the UBQ gene serving as an internal control. The primers used are listed in **Supplemental Table S2**.

To globally profile mRNA expressions in tobacco, the purified total RNAs (mentioned above) were sequenced via the platform of NovaSeq X Plus with the pair-end 150-nt mode. For the determination of gene expressions, the read trimming via Trimmomatic, the read alignment via STAR and the calculation of gene FPKM values were processed as described previously (Liu, MJ *et al*., 2018). The *N. benthamiana* draft genome sequence (v2.6.1) was retrieved from Sol Genomics Network (https://solgenomics.net).

The differentially expressed genes were identified via EdgeR analyses as described previously (Liu, MJ *et al*., 2018). The genes with |log2 fold-change| >=1 and FDRs <0.01 between the Pi-deficiency and -sufficiency conditions were retrieved for GO enrichment analyses. GO term enrichment analysis was performed with The Gene Ontology (GO) knowledgebase (http://geneontology.org)(Gene Ontology *et al*., 2023) to retrieve the gene count and the significance of enrichments in the indicated GO terms.

### Machine learning for the prediction of bioluminescent plants

The image datasets included three sets of biological replicate#1, #2, and #3, with plants grown under different Pi concentrations from Day 0, 11, 14 and 18. As indicated in **Fig. 6A**, the plants were categorized into different groups regarding the genetic backgrounds and Pi-deficiency treatments for the generation of models predicting either auto-luminescent plants and Pi-deficiency-induced signals in plants.

The prediction based on deep-learning methods can be divided into two parts. The first part involves image classification with two subtasks. Subtask 1 is to classify the auto-luminescent 35S plants from vector ones (yellow vs. gray in **Fig. 6A**). Subtask 2 is to classify the Pi-deficient plants versus Pi-sufficient ones of Sensor #25 and Sensor #31 (orange vs. blue in **Fig. 6A**). Several pre-trained classification models available in Torchvision or Keras were used including DenseNets (Huang *et al*., 2017), EfficientNets (Tan & Le, 2019), Inception (Szegedy *et al*., 2016), MobileNetV2 (Sandler *et al*., 2018), MobileNetV3-Large (Howard *et al*., 2019), ResNets (He *et al*., 2016), VGG-16 (Simonyan & Zisserman, 2015), VGG-19 (Simonyan & Zisserman, 2015) and Xception (Chollet, 2017) (see details of the tested models in **Supplemental Table S3**). The performances of these models were evaluated by several metrics, including accuracy, precision, recall, F1-score, and area under the ROC curve (AUC).

The second part of deep-learning-based predictions is the plant leaf area segmentation to evaluate the focus area of the classification models (i.e., the regions of the images that influence model performances). Grad-CAM, a technique to visualize important image regions influencing a deep learning model’s decision, was used to interpret the trained EfficientNets models (Selvaraju *et al*., 2017). We employ SegFormer, a modern and effective semantic segmentation framework (Xie *et al*., 2021), and YOLOv9, a widely-used model for image recognition and segmentation (Wang *et al*., 2024), to detect autoluminescent areas in plants. The purpose is to confirm the precise regions of interest in the trained EfficientNets models. SegFormer stands out due to its innovative Transformer-based encoder, which creates multiscale features without relying on positional encoding. This helps avoid performance issues when there are differences between training and testing image resolutions. It also uses a simple multi-layer perceptron decoder that merges information from different layers, combining local and global attention for strong results. YOLO, introduced by Redmon (2016), detects objects by splitting the image into grid cells and predicting object locations and classes. YOLO is known for its speed and accuracy in object detection, making it a strong candidate for identifying plant auto-luminescence.

The deep learning models used were based on Python 3.11, Pytorch 2.3.1, and TensorFlow 2.4.1. For all training cases, the entire images were cropped to 150 × 150, increasing the amount of data. Subsequently, the cropped images were resized to the specific size required for the particular models to train the models.

## RESULTS

### The generation of genetically encoded auto-luminescent tobacco biosensors that glow upon phosphate (Pi) deficiency

To employ fungal bioluminescent systems for reporting endogenous stress signals in plants, we genetically introduced fungal genes involved in luciferin synthesis and recycling including *NpgA* (phosphopantetheinyl transferase), *HispS* (hispidin synthase) and *H3H* (hispidin-3-hydroxylase), *CPH* (caffeoyl pyruvate hydrolase) as well as *Luz* (luciferase) into tobacco via agroinfiltration-mediated genetic transformation (**Fig. 1A**) (Khakhar *et al*., 2020; Mitiouchkina *et al*., 2020). The luciferin-synthesis and recycle-related genes including *NpgA, HispS, H3H* and *CPH* were driven by constitutive promoters (see Methods) whereas the luciferase (Luz) gene expression was controlled by the promoters of 35S (referred to as “35S”; a constitutive promoter as a positive control, referred to “35S”), “no promoter” (referred to as “Vector”; no promoter sequences as a negative control), and the phosphate-deficiency response promoter, TPSI1 (referred to as “Sensor”, being the sensor plant detecting Pi-deficiency) (Lin *et al*., 2016; Mitiouchkina *et al*., 2020) (**Fig. 1A**). The *TPSI1* gene encodes a long non-coding RNA-like mRNA in tomato and functions as a miRNA sponge to regulate gene expression and physiological response (Liu *et al*., 1997; Lin *et al*., 2016). Previous studies have reported that the tomato *TPSI1* gene is quickly induced across different plant stages and tissues including leaves (Liu *et al*., 1997). The *TPSI1* promoter (the 1.5-kb region upstream of the gene) has been experimentally validated to drive Pi deficiency-responsive gene expressions via the GUS reporter systems in tomato and tobacco (Lin *et al*., 2016). Thus, this *TPSI1* promoter was chosen to drive luciferase gene expression under phosphate deficiency here (**Fig. 1A**).

We focused on T2 homozygous tobacco plants and successfully isolated multiple independent and stable transgenic lines of Sensor, 35S and Vector plants. The Sensor and 35S plants, but not the Vector ones, showed auto-luminescence when grown in agar boxes **(Fig. 1B-D)**. These bioluminescent signals were autonomously emitted from plants and did not add the exogenous luciferin substrate and could be captured by a low-cost digital camera device (Nikon D5600 DSLR) (**Figs. 1,2A**). The auto-luminescence signal of the Sensor lines (#31 and #25) was predominantly higher under Pi deficiency condition (10 μM) than the sufficient one (500 μM) (**Fig. 1B**). In contrast, the 35S lines (#5 and #8) showed more comparable signals between sufficiency and deficient conditions whereas the Vector plants did not (**Fig. 1C,D**).

**Figure 2.**
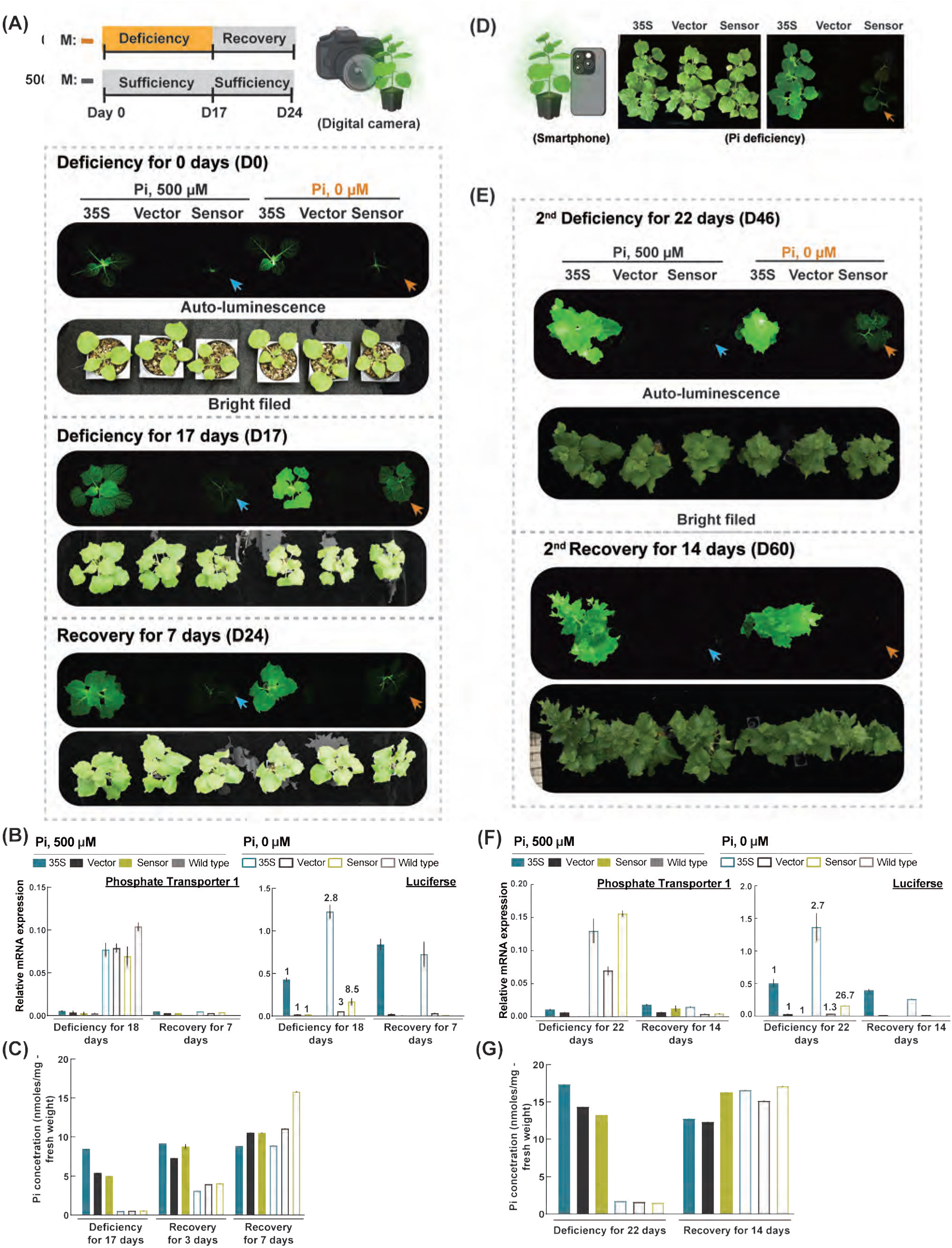
The responsiveness and robustness of bioluminescent sensor plants to Pi-deficiency. (A) As indicated in Fig. 1B, but for the auto-luminescent signals from 35S (#8), Vector (#1), and Sensor plants (#31) grown in vermiculite with nutrient solutions containing 500 uM phosphate (Pi) for ∼ 17 days (indicated as Day 0) and then subjected to Pi-deficiency (0 uM Pi, orange box), Pi-Sufficiency (500 uM Pi; gray boxes), and Pi-Recovery (500 uM Pi; gray boxes) treatments with indicated durations. (B) The quantitative real-time PCR analyses for the mRNA expressions of the fungal luciferase (*Luz*) gene and the tobacco Phosphate Transporter 1 gene for plants under Pi deficiency and sufficiency treatments at D17 as indicated in (A). The Ubiquitin 10 was used as an internal control for calculating the relative mRNA expressions. The normalized Luz expressions at the level of Pi deficiency/Pi sufficiency were shown. (C) The cellular Pi contents from the leaves of plants at D17 as indicated in (A). (D) As indicated in (A), but for the image of the plants at D17 captured by Iphone15 pro with 1-sec exposure and ISO 10,000. (E-G) As indicated in (A-C), but for the images (E), mRNA expressions (F) and Pi contents (G) of plants under the second round of the treatments with Pi deficiency for 22 days and then Pi resupply for 14 days. Error bars in (B,C,F,G) represent the SD values of the means of three technical replicates from one biological replicate. Consistent patterns were observed in two biological replicates as well as for the other set of the transgenic tobacco lines of 35S (#5), Vector (#7), and Sensor plants (#25) shown in **Supplemental Fig. S2**.

These results showed the success of generating independent Pi-deficiency responsive tobacco plants with strong auto-luminescence that did not depend on adding exogenous luciferin substrate and can be detectable by a consumer-level digital camera. In the following, one set of transgenic lines of 35S (#8), Vector (#1), and Sensor (#31) was included in the main figures whereas the other set was shown in supplemental figures.

### Sensor plants can responsively and robustly reflect the endogenous Pi status

The screening of Pi deficiency-responsive sensor plants with self-sustained auto-luminescent systems was conducted via agar box systems (**Fig. 1**). We further assessed the responsiveness of plants grown in vermiculite. Similar to the observation from agar boxes, the Sensor plants exhibited significantly stronger auto-luminescent signals when subjected to phosphate deficiency (0 μM) whereas these signals were decreased during recovery conditions (i.e., phosphate-sufficient treatment; 500 μM) (orange and blue arrows in **Fig. 2A; Supplemental Video 1**). In contrast, the 35S plants showed signals in both Pi deficiency and sufficiency conditions while the Vector plants did not (**Fig. 2A**). The mRNA abundance of the luciferase gene in Sensor plants showed a ∼8.5-fold increase under phosphate deficiency (open yellow bars; right panel in **Fig. 2B**), which was much higher compared to ∼2-3-fold increase in 35S and Vector plants, supporting the Pi-induced transcriptional response driven by *TPS1* promoter (Liu *et al*., 1997; Lin *et al*., 2016). The mRNA expression of Phosphate Transporter 1, a known Pi-deficient induced gene (Chiou *et al*., 2001; Chien *et al*., 2022), was induced across transgenic plants as well as the wild-type tobacco (left panel in **Fig. 2B**). In addition, the cellular Pi contents in Sensor plants were low under deficiency conditions and increased during the recovery treatment (**Fig. 2C**). These observations showed that the bioluminescent signals in Sensor plants were responsive to the dynamic change of external Pi abundances. The results further demonstrated that the bioluminescent signals, as an easy-captured readout, could reflect the *in planta* Pi status and gene expressions across different transgenic plants including Sensor plants.

To assess the robustness of Sensors to Pi deficiency, we conducted two rounds of Pi-deficiency treatments in a row by putting the first-round recovered plants into Pi deficiency and then sufficient conditions. The analyses of auto-luminescence signals, mRNA expressions of the Luz gene, and cellular Pi contents reflect the impacts of the second-round Pi-deficiency treatments (**Fig. 2E-G**), showing the repeatable responsiveness of Sensor plants to external Pi dynamics. Similar patterns were also observed in other independent lines **Supplemental Fig. S2**. Furthermore, we explored the extent to the auto-luminescent signals of Sensor plants could be detected with low-cost devices. We found that these auto-luminescent signals were detectable not only using a digital consumer-level camera (**Fig. 2A**) but also by the camera of smartphones such as iPhones (**Fig. 2D**), and even by the naked eye in darkness. This showed the potential of using Sensor plants in both lab research and field application.

To assess the potential of the Sensor plants as a readout of stress status before physiological damages, we examined the dynamic changes of auto-luminescent signals and plant morphology during Pi deficiency. The Pi-deficient induced auto-luminescence signals were observed at ∼10 days whereas the plants became smaller and pale green at ∼14 –17 days (**Supplemental Fig. S1; Supplemental Video 1**), similar to the phenotypes of the Pi deficiency in tobacco observed previously (Lin *et al*., 2016). This observation reflected the transcriptional regulation of genes occurring before the physiological damage in plants.

Together with previous observations for plants grown in agar plants (**Fig. 1**), these findings showed the responsiveness and robustness and of auto-luminescent biosensors in detecting endogenous Pi status in plants. In addition, the Sensor plants provide high-sensitivity signals recorded by low-cost devices before the morphological changes.

### The Pi-deficiency-induced gene expressions of plants with genetically encoded bioluminescence systems

Previous studies reported that introducing genetically encoded bioluminescence systems into tobacco did not lead to significant phenotypic differences including plant growth and yield, except the plant height and the leaf necrosis under strong illumination (Mitiouchkina *et al*., 2020; Shakhova *et al*., 2024). Since this system was coupled with Pi-induced promoters to detect plant endogenous Pi stresses here, we further assessed its impact on Pi deficiency response in plants by globally profiling gene expressions via mRNA-sequencing analyses. The correlation analyses showed the global expression profiles among three distinct transgenic plants and wild-type tobacco were all highly similar with the Spearman’s correlations >0.92 (**Fig. 3A**). We noted that, despite of the different genetic backgrounds, the Pi-deficiency-treated samples were grouped and separated from the Pi-sufficiency-treated ones (**Fig. 3A**). In line with the observation, the Principle Component Analyses based on global gene expressions also showed that the phosphate treatments, but not the genetic background, was the primary factor in clustering samples (PC1 in **Fig. 3B**).

**Figure 3.**
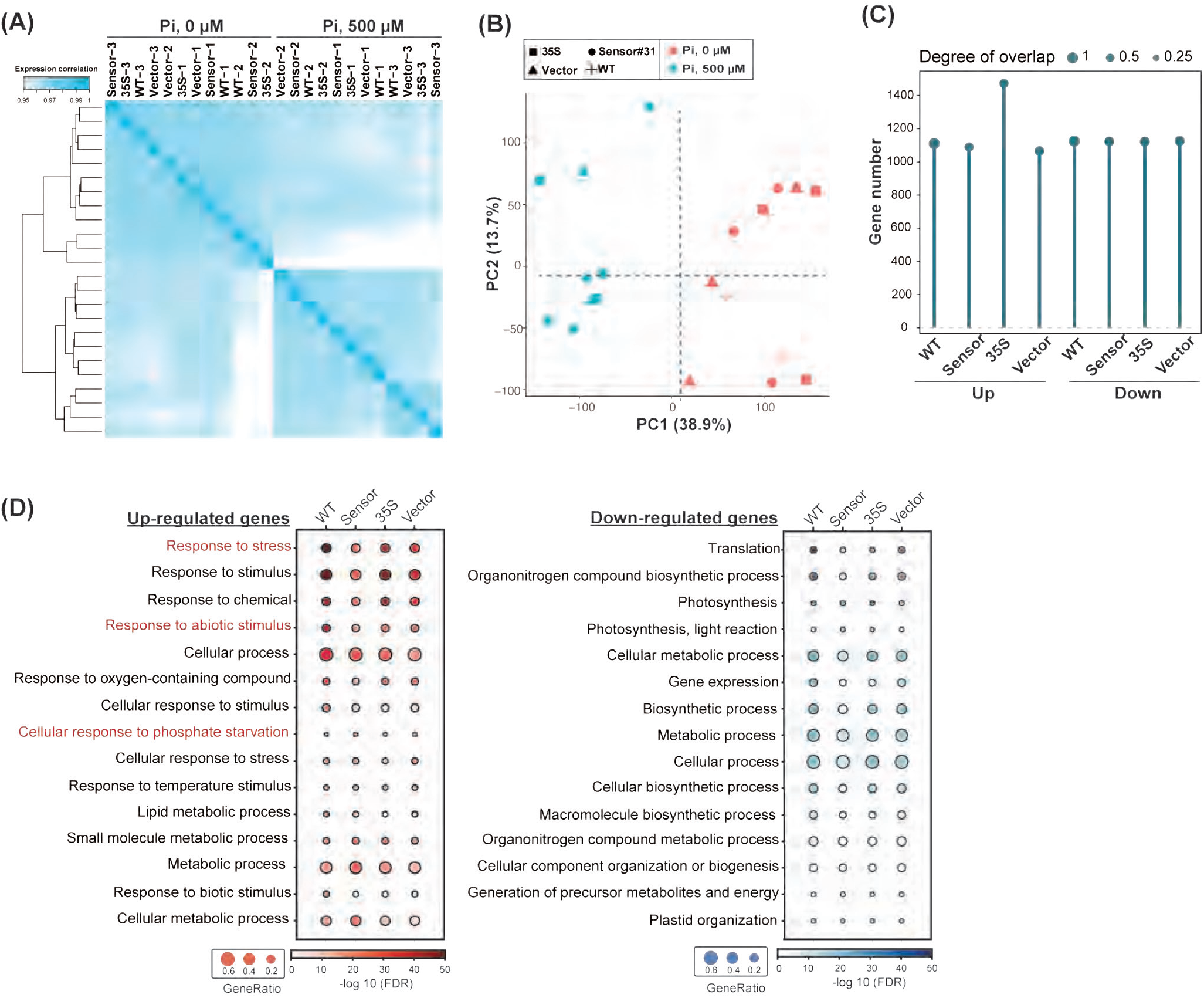
Global gene expression profiles of sensor plants upon Pi deficiency. (A) The Spearman’s correlation analyses of global gene expressions for plants indicated in Fig. 2A and wild-type (WT) tobacco with the treatments of Pi deficiency (0 uM) and sufficiency (500 uM). Shown for the samples collected from 3 biological replicates. (B) As indicated in (A), but show the Principle component analyses for WT and different transgenic plants under Pi deficiency (0 uM; pink circles) and sufficiency (500 uM; cyan circles). (C) The number of the differentially expressed genes upon Pi-deficency treatment (|log2 fold-change| >=1 and FDRs <0.01) and the overlap of genes (proportion) identified from 35S, Vector and Sensor plants relative to those identified from WT. (D) Gene Ontology analyses for the Pi-deficiency regulated genes (as indicated in (C)). Shown the adjusted *p-*values and gene counts (%) across different transgenic lines and WT plants for the top15 Biological process-related terms with the lowest adjusted *p-*values in WT plants.

In addition, more than 63% of Pi-deficiency-regulated genes in transgenic lines overlapped with those identified from wild-type plants including the known Pi-deficiency induced genes of *PHO1* and *SPX* gene family *(SPX1-SPX4)* (**Fig. 3C; Supplemental Fig. S3B**) (Ribot *et al*., 2008; Liu, N *et al*., 2018). The Gene Ontology analyses found that the significantly induced genes upon Pi deficiency were enriched in the GO terms of Biological process related to stress, abiotic stimulus and phosphate starvation. This observation was found in both transgenic and wild-type tobacco (**Fig. 3D).** Additionally, the transgenic and wild-type tobacco plants exhibit similar patterns in the GO terms for Biological Processes and Molecular Functions (**Supplemental Fig. S3A)**. Lastly, the dynamics of the morphological changes among 3 transgenic lines were similar when subjected to Pi deficiency. The three transgenic lines all turned into pale green and reduced growth at ∼ 14-17 days after Pi deficiency treatments (**Supplemental Fig. S1**).

Together, the analyses of global and Pi-deficiency-induced gene expression profiles as well as the morphological changes showed the similarity of Pi deficiency responses between wild-type tobacco and the transgenic plants (including 35S, Vector and Sensor plants) with genetically encoded bioluminescence systems. This suggests that introducing fungal bioluminescence systems into plants did not significantly impact their responses to Pi deficiency.

### Sensor plants are specific and sensitive to a wide range of phosphate deficiency conditions

Soils contain multiple essential macronutrients including phosphors as well as nitrogen and potassium. We next asked, in addition to phosphors, whether the sensor plants are responsive to other nutrients. The plants were grown in vermiculite and cultured with nutrient solutions in which specific nutrients were depleted (**Fig. 4A**). As expected, compared to the full nutrient solution (i.e., the solution with nitrate, phosphate and potassium; See methods), the Sensor plants showed higher signals when treated with the Pi-depleted solution (orange arrow, **Fig. 4B**). The lower cellular Pi concentration and the higher gene expressions of the *Luz* and *Phosphate Transporter 1* genes reflected the higher luminescent signals and Pi-deficiency treatment (**Fig. 4C**; **Supplemental Fig. S4** for other independent lines). We noticed that a minor increase of auto-luminescent signals and the mRNA abundances of the *Luz* gene in Sensor plants under nitrogen and potassium deficiency conditions (**Fig. 4B,C** and **Supplemental Fig. S4B,C**). The cellular Pi contents were comparable for plants grown under nitrogen/potassium deficiency and full-nutrient conditions (**Fig. 4D**). These results indicated that *TPSI1* promoter was minorly responsive to nitrogen/ potassium deficiency. Together, our observations indicated that the Sensor plants majorly responded to Pi deficiency.

**Figure 4.**
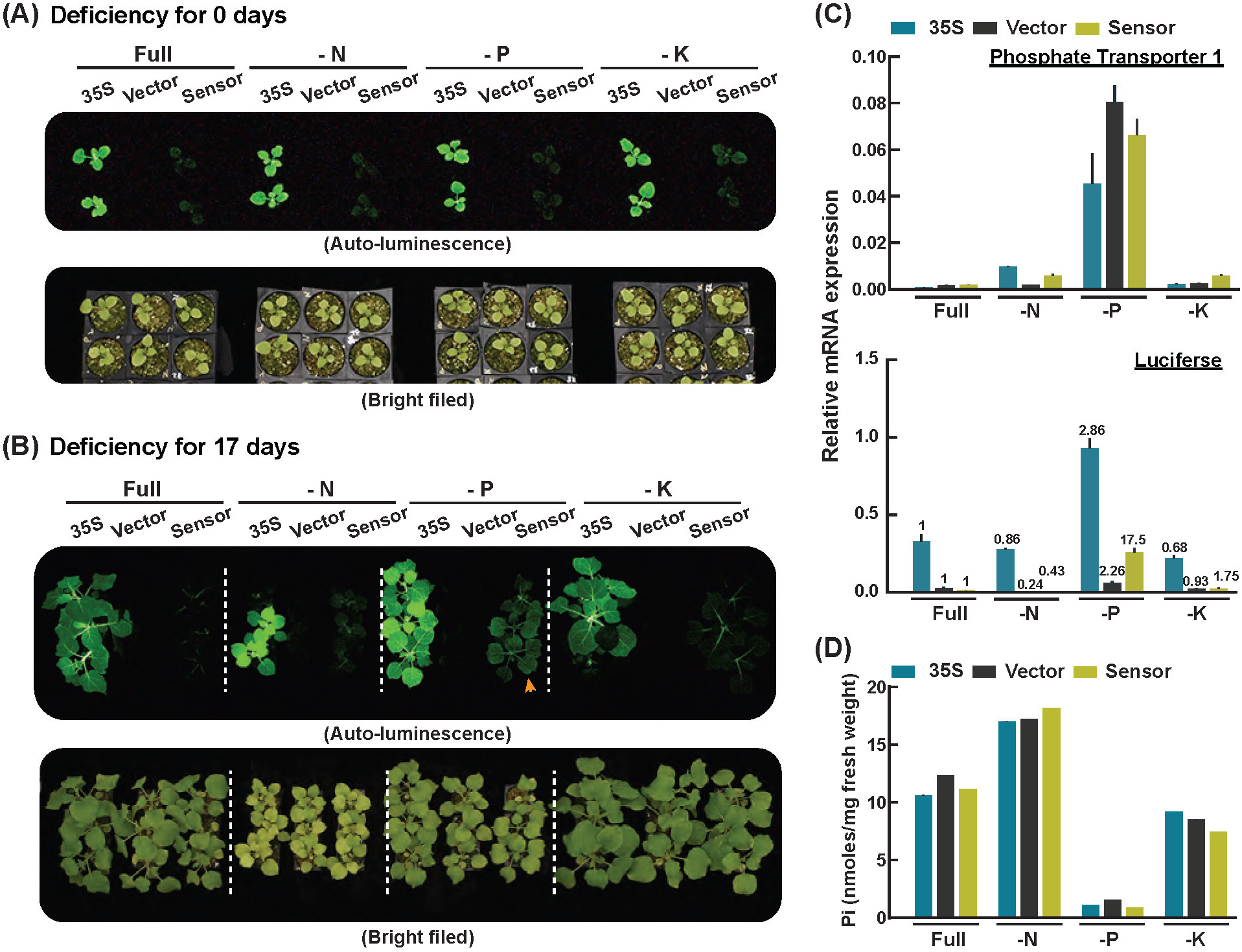
The nutrient-specificity in inducing the auto-luminescence of sensor plants. (A) The auto-luminescent signals of the 35S (#8), Vector (#1), and Sensor plants (#31) grown in vermiculite for ∼ 17 days (indicated as Day 0) and subject to the withdrawal of the nitrate (-N), phosphate (-P) and potassium (-K), respectively, and without the nutrient withdrawal (Full). (B) As indicated in (A), but for transgenic tobacco plants under nutrient deficiency treatments for 18 days. (C,D) As indicated in **Fig. 2B,C**, but for the mRNA expressions of indicated genes (C) and the cellular Pi contents (D) from the leaves of the plants under different nutrient deficiencies as indicated in (B). The normalized Luz expressions at the level of Pi deficiency/Pi sufficiency were shown. The results for the other set of the transgenic tobacco lines of 35S (#5), Vector (#7), and Sensor plants (#25) were shown in **Supplemental Fig. S4**.

The findings so far were based on sensor plants grown in the laboratory-used plant medium including agars and vermiculite (**Figs. 1-4**). In addition, the observations of Pi-deficiency triggered response were based on one arbitrary concentration (0 μM in vermiculite and 10 μM in agar boxes). To further explore the responsiveness and sensitivity of Sensor plants and the impact of culture medium in plant stress responses, the plants were grown in soils (with limited Pi contents, **Supplemental Table. S1**) and watered with nutrient solutions with five different Pi concentrations ranging from 0 to 500 μM (**Fig. 5A**; see **Supplemental Fig. S5** for the other replicate). The Pi concentration lower than 50 μM could induce higher luminescent signals, an observation in both #31 and #25 sensor lines. We noticed that compared to the #25 sensor line, the #31 sensor lines exhibited more predominant Pi-deficient induced luminescence, which can be clearly observed on day 10. This was probably due to the higher basal auto-luminescent signals of #31 lines, as detected at day 0, that facilitated the visualization of the signal difference among different treatments (**Fig. 5A; Supplemental Fig. S5A**). In addition, the induced mRNA expressions of *Luz* and *Phosphate Transporter 1* genes and the lower cellular Pi contents were observed across the conditions with ≤ 50 μM Pi, correlating to the observed luminescence signals, in sensor plants (**Fig. 5B,C**).

**Figure 5.**
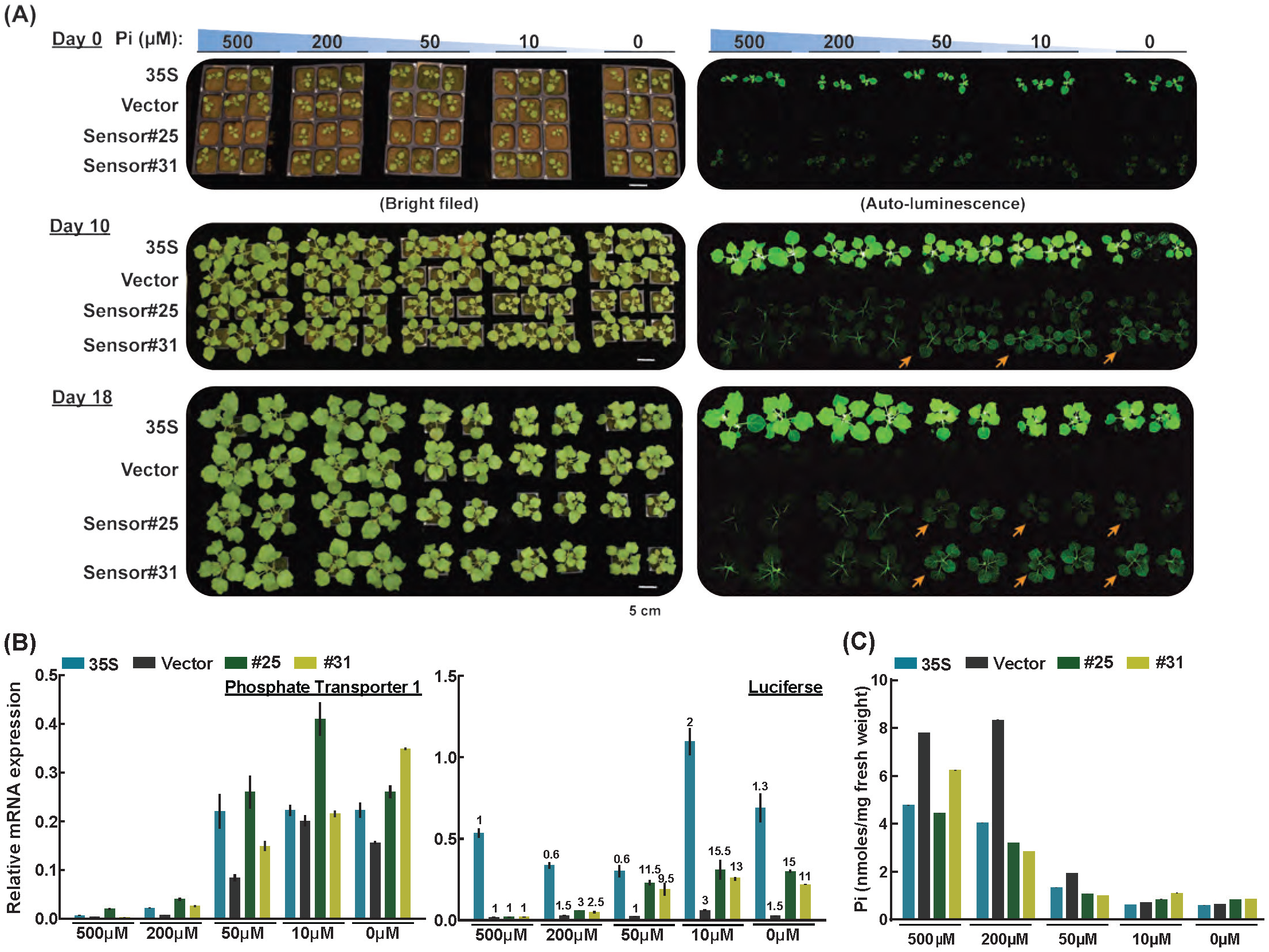
The sensitivity of sensor plants in response to Pi deficiency. (A) As indicated in Fig. 1B, but for the auto-luminescent signals from 35S (#8), Vector (#1), and Sensor plants (#31 and #25) grown in Pi-deficiency soils with nutrient solutions containing 500 uM phosphate (Pi) for ∼ 17 days (indicated as Day 0) and then subjected to the treatments with nutrient solutions containing five different Pi concentrations. (B,C) As indicated in Fig. 2B**,C**, but for the mRNA expressions of indicated genes (B) and the cellular Pi concentrations (C) from the leaves of the plants under different nutrient deficiencies as indicated in (A) at Day 18. The normalized Luz expressions at the level of Pi deficiency/Pi sufficiency were shown.

Together, these results showed that the responsiveness of sensor plants was observed across different growth conditions and induced at a threshold of Pi deficiency concentration. In addition, Sensor plants can majorly respond to Pi deficiency but not nitrogen/potassium deficiency.

### Deep-learning-based image analyses enable predicting Pi deficiency in plants

A quantitative imaging system, coupled with machine-learning techniques, capable of remotely and longitudinally recording luminescence and reporting stress signals is essential for field-scale plant sensing. The system’s ability to distinguish luminescence signals from background levels is crucial and could be achieved via machine-learning-based object classification techniques to develop a model for classifying plants such as sensor plants under control conditions and subjected to Pi deficiency treatments.

By employing the imaging datasets collected from the sensor plants using Nikon D5600 DSLR camera (**Fig. 5A**), we used deep-learning-based object classification techniques, including different Convolutional Neural Network (CNN)-based algorisms, to build a model to classify the sensor plants under Pi deficiency and under Pi sufficient conditions. We focused on the sensor plants under Pi deficiency (i.e., 0, 10, and 50 μM Pi) and those under Pi-sufficiency (i.e., 200 and 500 μM Pi) conditions for 11-, 14- and 18-day treatments (orange vs. blue in **Fig. 6A**; See **Methods**). We found that, among different tested algorisms, multiple classification methods all performed well, achieving the mean of AUROC values >=0.9, in distinguishing the Pi-deficient and Pi-sufficient sensor plants (**Fig. 6B; Supplemental Fig. S6B**).

**Figure 6.**
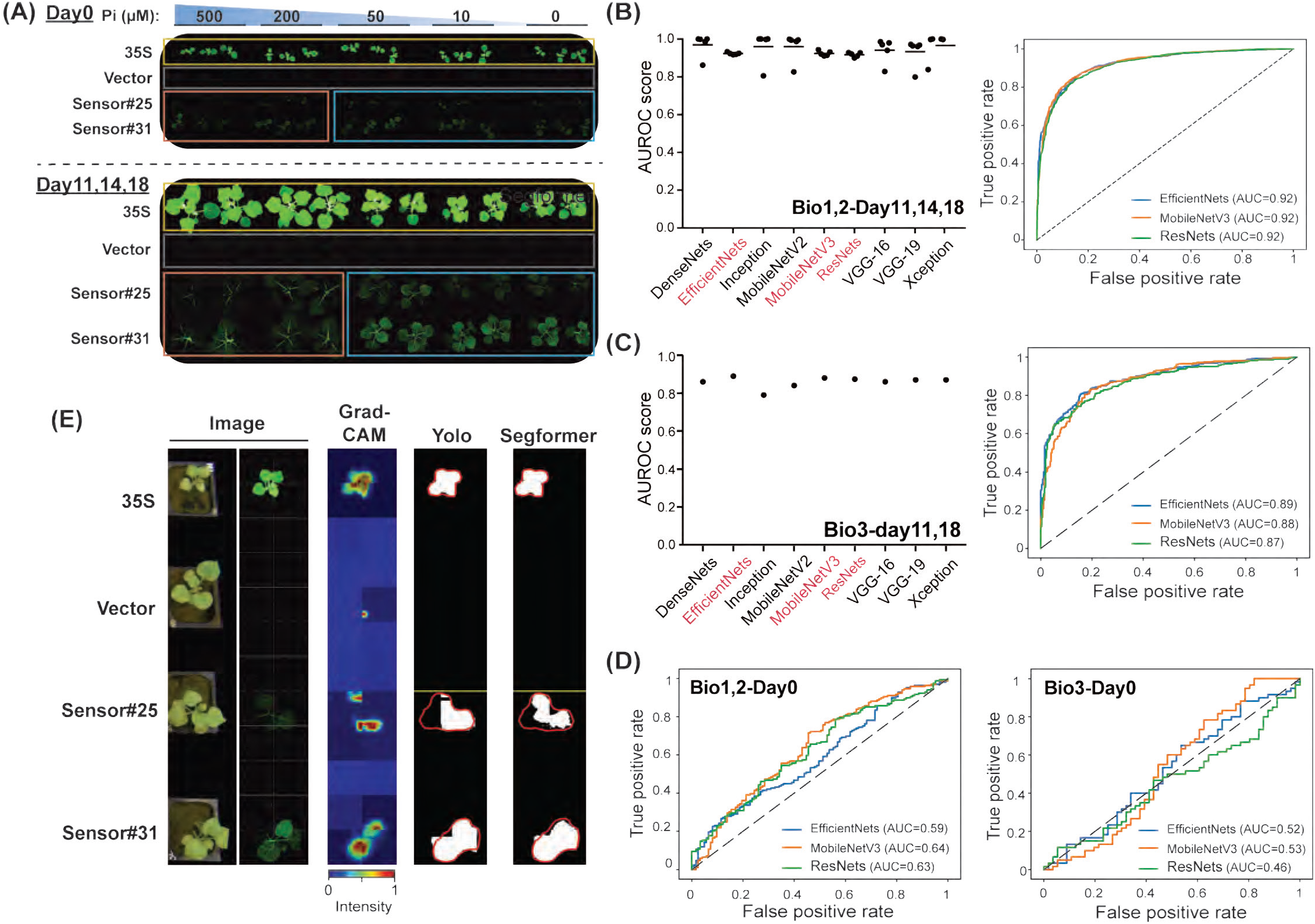
The integration of imaging analyses and computational modeling enables to classify sensor plants under Pi deficiency conditions. (A) Illustration of the image datasets for generating models for predicting Pi-deficiency-induced auto-luminescent plants. As depicted in Fig. 5A, plants are grown under various conditions and durations. Yellow and gray boxes: the area of images used for building and testing the models of detecting luminescent signals (subtask1; see results in **Supplemental Fig. S6A**). Orange and blue boxes: the area of images used for building and testing Pi-deficiency prediction models (subtask2). (B) The prediction performance of subtask2 (shown in AUROC values) for various classification models using the datasets collected from Day11,14 and 18 of bioreplicate#1 and #2 to distinguish the plants under Pi sufficient (orange) and deficient conditions (blue) as indicated in (A). Left panel: the mean and its individual AUROC values from five-fold cross-validation. Right panel: the AUROC curves of the mean from five-fold cross-validation. See **Supplemental Fig. S6B** for other indications of model performances. (C) The prediction performance of using the models built from (B) to classify the plants (orange vs. blue; as indicated in (A)) based on the images collected from Day11 and 18 of bioreplicate#3. (D) The prediction performance of the EfficientNets, MobileNetV3-Large, and ResNets built from (B) in classifying plants (orange vs. blue; as indicated in (A)) based on the images collected from Day0 of bioreplicate#1 and #2 (left) and the Day0 of bioreplicate#3 (right). (E) The image showed the plants under 0 uM Pi for 18 days (as indicated in (A)), which was collected from bioreplicate#3. The Grad-CAM heatmap displayed the focused area of EfficientNets whereas the warm color indicates a greater intensity of focus. The segmentation results for the YOLOv9 and SegFormer models were depicted as white masks, and the red line outlines the ground truth area.

To assess the robustness of the build Pi-deficiency prediction models, we further asked to what extent the build models can predict the Pi status of the sensor plants based the images collected from the independent bioreplicate. Most models achieved AUC values ranging from 0.79 to 0.88 (**Fig. 6C**). These results showed the robustness of the build models that based on Pi deficiency-induced luminescent signals can predict the plants under Pi deficiency. Notably, the methods of EfficientNets, ResNets and MobileNetV3-Large showed high consistency with minimal variances throughout the model-building process (i.e., the five-fold validation; **Fig. 6B**) as well as achieved the best performance in assessing the robustness of methods (**Fig. 6C**). In addition, when applying these generated models to predict the sensors plants based on the images of Day-0 sensor plants, the model performances dropped with the AUCROC values of 0.46∼0.64 (**Fig. 6D**), reflecting the comparable and low luminescent signals of Day-0 sensor plants between Pi-deficiency and -sufficiency conditions (**Fig. 6A**). We would like to note that this deep-learning-based analytical pipeline generated a model that can well differentiate auto-luminescent plants of 35S lines from the vector ones (yellow vs. gray in **Fig. 6A**; **Supplemental Fig. S6A**). This showed the capability of the analytic pipeline and the generated model to detect autoluminescent expression in plants.

To visualize the important features of an image for model prediction, we conducted Grad-CAM analyses to highlight the crucial leaf regions focused on by the EfficientNets model. The Grad-CAM heatmap clearly showed that the regions with greater intensity of model focus tend to have higher auto-luminescent signals in plants (**Fig. 6E**). The YOLOv9 and SegFormer, two well-established image segmentation models (Xie *et al*., 2021; Wang *et al*., 2024), performed well in segmenting the auto-luminescent areas from 35S plants (white masks in **Fig. 6E)**, with the performances of mAP50=0.936 in YOLOv9, and the dice coefficient=0.96 in the SegFormer (**Supplemental Table S4)**. This showed that both models could detect luminescent signals of leaf regions from 35S plants and the detected regions were aligned with the focus area of Grad-CAM (**Fig. 6E**). Focusing on sensor plants (#25 and #31), the luminescent regions were highlighted in Grad-CAM heatmap as well as identified by YOLOv9 and SegFormer (**Fig. 6E**). Nevertheless, the regions with dim luminescence such as the left side of sensor #25 were not identified, showing the limitation of the current models. The results indicate that EfficientNets focused on the luminescent signals and can predict Pi-induced bioluminescent plants.

Together, we developed a deep-learning-based image analysis pipeline capable of capturing luminescence signals from the aerial parts of living plants and created a predictive model that infers stress status based on luminescent signals and provides predictions about soil stress.

### Application of Pi-deficiency-monitoring biosensors to different crops via grafting

Our findings so far demonstrate the successful generation of a Pi-deficiency-sensing plant biosensor with self-sustained bioluminescence responsive to endogenous Pi status in tobacco (**Fig. 2,5**). Facing diverse crops, how can the technique of stress-induced auto-luminescent biosensors be applied to different crops for monitoring their Pi-deficiency stress? One bottleneck is to genetically introduce fungal bioluminescence systems into target crops, which is a time- and effort-consuming process. To meet this challenge, employing the grafting technique, we asked whether the shoot part of tobacco sensor plants, when grafted onto the target crops such as tomato, responds to Pi deficiency (**Fig. 7A,B**)

**Figure 7.**
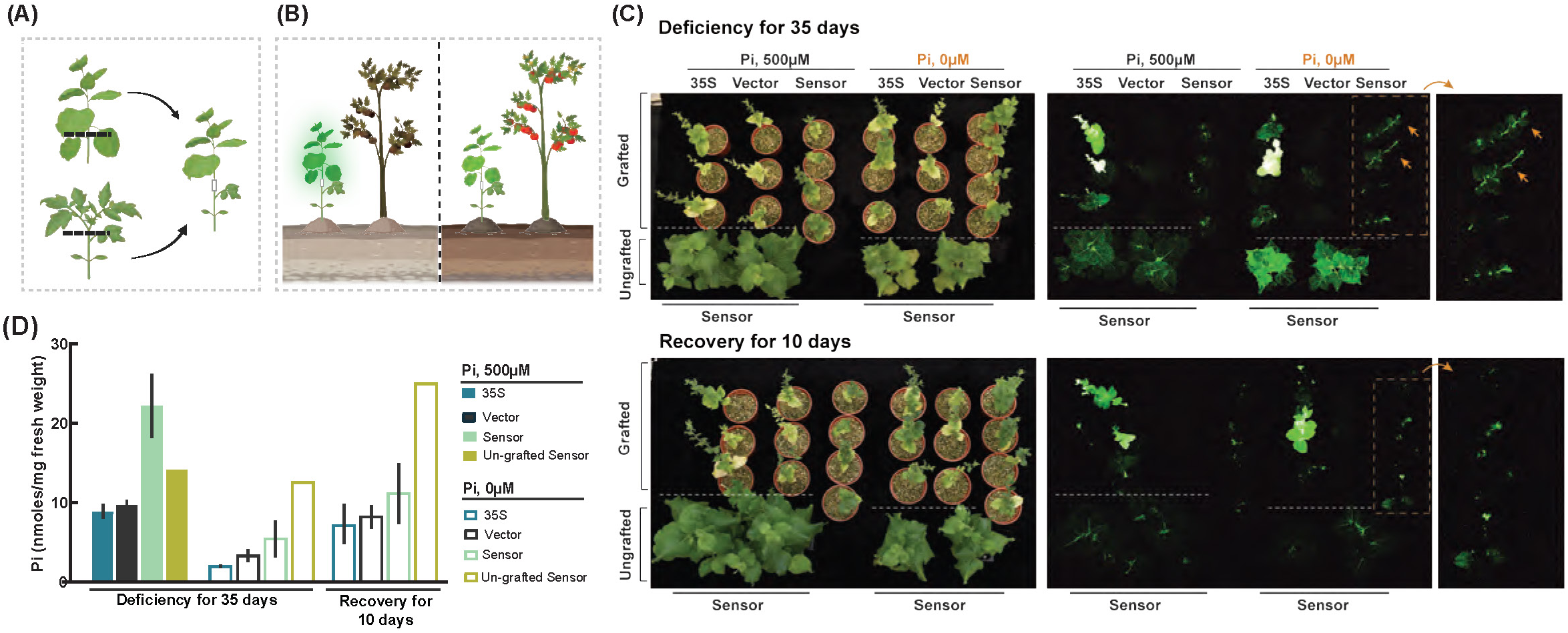
Sensor plants grafted onto different crop species responded to Pi-deficiency. (A,B) Schematic of grafting Sensor plants onto target crops (A) for monitoring their Pi deficiency response under Pi deficiency and sufficiency conditions (left and right panels in (B)). (C) The auto-luminescent signals of the 35S (#8), Vector (#1), and Sensor shoots (#31) that were grafted onto tomatoes since grafting (indicated as Day 0) and subjected to Pi deficiency (0 uM) for 35 days and then Pi resupply for 10 days. Dashed orange box: the area with the zoom-in view. (D) As indicated in (C), but for the cellular Pi concentrations (C) from the leaves of the grafted plants. (See **Supplemental Fig. S7** for the results of chili pepper).

By grafting the tobacco sensor shoot onto the tomato seedling, we found that the sensor-grafted tomato plants exhibited higher auto-luminescent signals upon Pi deficiency than under Pi sufficiency condition (orange arrows in top panels; **Fig. 7C**). This induced auto-luminescent signal was decreased when subjected to the recovery condition (bottom panels; **Fig. 7C**). This observation was also found for the tobacco sensor plants without grafting (as a positive control) that showed induced and reduced auto-luminescence under Pi deficiency condition and subjected to the recovered condition (**Fig. 7C**). In addition, the Pi-treatment-dependent auto-luminescence signals were correlated with the cellular Pi contents in the grafted sensor leaves (**Fig. 7D**).

To further explore the potential of applying sensor plants to detect Pi deficiency of diverse crop species, we also grafted sensor plants onto another crop, the chili peppers. Similar to the observation in sensor-grafted tomato plants, we found that the tobacco sensor part of the grafted chili peppers was responsive to Pi status, showing different magnitudes of auto-luminescence under Pi deficiency and sufficiency conditions (**Supplemental Fig. S7A**)

Together, these results showed that the tobacco sensors, when grafted onto another crop species, could respond to Pi-deficiency treatment, inferring the endogenous Pi-deficiency stress of the target crop. With the integration of the grafting technique and the ability of tobacco to form grafts with a diverse range of angiosperms (Notaguchi *et al*., 2020), our findings suggest the wide-ranging application of tobacco sensor plants to monitor the Pi deficiency stresses of diverse crops.

## DISCUSSION

Our study generated a Pi-deficiency sensing-reporting plant biosensor with self-sustained bioluminescence that was responsive to the endogenous Pi status and detected via low-cost devices and deep-learning based models. We first coupled the fungal auto-luminescent systems (including the self-sustained luciferin synthesis system and the luciferase gene) and the plant Pi-deficiency responsive promoter to transgenically generate the sensor plants (**Fig. 1**). The sensor plants showed autonomous bioluminescence, which can be observed by the naked eye and captured by consumer-grade cameras/smartphones, and the light emission was induced upon Pi deficiency (**Fig. 2,5**). The introduction of fungal luminescent systems into plants did not impact the global gene expression and the Pi-deficiency response (**Fig. 3**). The auto-bioluminescent sensor plants showed the inducibility and specificity to Pi deficiency but not other nutrient deficiency, which were observed when grown both in agar/vermiculite medium and soil (**Fig. 1-5**). The deep-learning based model based on the bioluminescent image collected can efficiently predict the sensor plants under Pi deficiency conditions (**Fig. 6**). Lastly, the grafted sensor plants (with sensors in the scions and target crop in the rootstocks) responded to Pi deficiency, (**Fig. 7**). Together, these finding showed, complementary to the mechanical and other spectrum-image-related biosensors (de Ruijter *et al*., 2003; Saric *et al*., 2022), our self-sustained bioluminescent sensor provided a high-inducibility and high-sensitivity light system that can detect *in vivo* and real-time stress status and be captured via non-invasive and low-cost devices. The integration of deep-learning and grafting techniques provides the potential of applying the sensors to various crops and establishing a monitoring-reporting system of plant stresses in agricultural fields.

Our results of CNN-based bioluminescent prediction models including EfficientNets, MobileNetV3-Large, and ResNets exhibited the lowest standard deviation of AUC performance and the highest and robust AUC performance in the datasets of an independent bioreplicate (**Fig. 6B,C**). The trained models such as EfficientNets was capable of identifying the partial area of the leaf with a luminescence signal, enabling the detection of Pi deficiency within a larger area (**Fig. 6E**). Besides, MobileNetV3-Large and EfficientNets had relatively small parameter counts and model sizes (**Supplemental Table S3**). Thus, from a practical standpoint, MobileNetV3-Large and EfficientNets will be more suitable for deployment in the future. Additionally, our CNN-based models were effective in distinguishing luminescence signals between the Pi-sufficient (200 and 500 uM) and Pi-sufficient (0, 10 and 50 uM) groups (orange and blue groups in **Fig. 6A**). In the future, it will be valuable to develop a multi-classification model to discern the different levels of Pi deficiency among plants, enabling more precise determination of the *in vivo* Pi-deficiency status in plants.

The intensity of light emission plays a critical role in facilitating light capturing via low-cost devices. Our bio-luminescent sensors showed increased light emission upon Pi deficiency whereas the brightness of luminescence was lower than the 35S lines (**Fig. 1**). This observation highlighted the relatively low luminescent intensity and the need of further optimization of its sensitivity and the light emission magnitude. A recent study targeting on genes in the fungal bioluminescence pathway and employing consensus mutagenesis as well as screened orthologous genes successfully improved the thermostability and catalytic activities of enzymes. This optimized bioluminescence pathway enhanced one to two orders of luminescence magnitudes across various heterologous hosts, including plants (Shakhova *et al*., 2024), representing a genetic engineering strategy of optimizing bioluminescent systems. Alternatively, the mRNA stability, codon optimization and the translational enhancer that affects protein synthesis efficiency can achieve higher expression of genes in the bioluminescence pathway as well as the accumulation of luciferin substrates (Chappell *et al*., 2015; Huang *et al*., 2021; Kim *et al*., 2022). It will be worth to adapt these genetic engineering methodologies to generate the next version of Pi-deficient responsive plant biosensor with better brightness. Nevertheless, when coupling the optimized bioluminescent systems and the plant Pi-deficiency responsive promoter (that drives *Luz* gene expression), whether bioluminescent signals can timely and precisely reflect the dynamic changes of endogenous stress status will need to be assessed.

In addition to the sensitivity of the biosensors (i.e., detectable reporter signals), the inducibility and specificity are also the key characteristics of the stress-monitoring biosensor. The Pi-deficiency responsive *TPSI1* promoter used in this study drove *Luz* gene expression and triggered auto-luminescence under Pi deficiency (**Figs. 1-3**). The *TPSI1* had a known motif site of P1BS, which is conserved in Pi-deficient response genes across different plant species (Bustos *et al*., 2010; Lin *et al*., 2016; Araceli *et al*., 2017) and showed an at least ∼ 8-fold increase of LUZ expression (**Figs. 2-5**). Studies showed that the number of motif sites as well as their flanking sequences within a Pi-induced promoter can affect the magnitudes of the Pi-deficient responsive promoter strength (Araceli *et al*., 2017). In line with this, global and bioinformatics analyses showed that the *cis*-regulatory elements work in a context- and cooperative-dependent manner to tune the quantitative strength of gene activation and repression (de Boer & Taipale, 2023; Kim & Wysocka, 2023). These findings indicate the potential of editing the *TPSI1* or other Pi-deficient related promoters to improve the inducibility of the stress-monitoring biosensor. In addition, our results showed a slight increase of auto-luminescent signals upon nitrogen/potassium deficiency (**Fig. 4**). One possibility is the crosstalk between stresses that different abiotic stresses can regulate the expressions of partially overlapping sets of genes. Indeed, consistent with the observation of nitrogen/potassium-phosphate interactions (Medici *et al*., 2019; Wang *et al*., 2021), the Phosphate Transporter 1 expression was minorly induced upon nitrogen deficiency (**Fig. 4C**). How and by which *cis*-elements that lead the *TPSI1* promoter responsive to nitrogen/potassium deficiency remains to be explored. Together, the inducibility and the specificity of a biosensor can be regulated at least by the promoter strength and *cis*-element contexts. Our results showed that the Pi-deficient *cis*-regulatory code remains to be comprehensively elucidated in order to govern when and how much each gene is transcripted and optimize the biosensor activity. The combination of machine learning and massively parallel assays employing synthetic DNA will be a strategy to not only crack the *cis*-regulatory code but also provide information of de novo designed sequences to facilitate the development of high-specificity and high-inducibility biosensors.

The introduction of fungal bioluminescent systems into plants eliminates the need to add exogenous *luciferin* substrate and turns plants themselves into auto-glowing ones, in which the light emission can be captured via low-cost cameras and analyzed via DL-based imaging analyses. Coupling with stress-induced LUZ expression systems, the generated plant biosensors are responsive to abiotic stress in soil and report the stress levels *in planta*. Together, the plant biosensor-supported and real-time imaging system sheds light to quantitatively record luminescent signals from living plant aerial parts and infer the *in vivo* stress status in agriculture.

## ACKNOWLEDGMENTS

We thank the core facilities of Transgenic Plant Laboratory and AS-BCST Bioinformatics Core for agrobacteria-infiltration in tobaccos and high-performance computing services, Dr. Shu-I Lin and Dr. Tzyy-Jen Chiou for sharing the TPSI1 promoter construct and the knowledge in performing Pi deficiency treatment in plants and Dr. Kuo-Chen Yeh and Dr. Hsin-Hung Yeh for intellectual input in conceiving the research. This research was financially supported by grants of the RSF 24-74-10087 to A.-S. M., the MRC Laboratory of Medical Sciences (UKRI MC-A658-5QEA0) to K.-S. S, and AS-TP-110-M07 and NSTC 113-2326-B-001 to Ming-Jung Liu.

## AUTHOR CONTRIBUTION

C.-W C., performed the bioinformatics and experimental analyses and image dataset collection. Y.-F. C. and Y.-R. L. performed the experimental analyses. L.-J. C., H.-J. L., H.-J. H. performed image and prediction analyses. T.-Y. L. performed image and prediction analyses and wrote the paper. A.-S. M. and K.-S. S. created the starting luminescence plasmids. C.-H. H. conceived/designed the research. M.-J. L. conceived/designed the research, performed sequencing analyses, and wrote the paper.

## DATA AVAILABILITY

All raw and processed sequencing data generated in this study have been submitted to the NCBI Gene Expression Omnibus (GEO; https://www.ncbi.nlm.nih.gov/geo/) under accession number GSE277579.

## SUPPLEMENTAL DATA

***Supplemental Figure S1. The Pi-deficiency-induced auto-luminescent signals and morphological changes of the sensor plants with Pi-deficiency-induced bioluminescent systems.***

As indicated in **Fig. 2, and for** the auto-luminescent signals and the bright-filed images of 35S (#8), Vector (#1), and Sensor plants (#31) grown in vermiculite with nutrient solutions containing 500 uM and 0 uM phosphate (Pi) for the indicated days.

***Supplemental Figure S2. The Pi-deficiency-induced responsiveness and robustness of the bioluminescent sensor plants***

(A-F) As indicated in **Fig. 2**, but for the luminescent signals under 1^st^ and 2^nd^ Pi-deficiency and recovery conditions (A,D), the mRNA expressions of phosphate transporter 1 and Luz genes (B,E), and cellular Pi contents (C,F) for the other independent lines of 35S (#5), Vector (#7) and sensor plants (#25).

***Supplemental Figure S3. The enrichments of the gene-ontology terms in sensor plants upon Pi deficiency treatment***

(A) Gene Ontology analyses for the Pi-deficiency-induced up-/down-regulated genes (|log2 fold-change| >=1 and FDRs <0.01). Show the adjusted *p-*values and gene counts (%) across different transgenic lines and wild-type (WT) tobacco plants for the top30 Biological process/Molecular function-related terms with the lowest adjusted *p-*values in WT plants. (B) Differential expression values (shown in log2 Fold-changes (FCs)) by contrasting Pi-deficiency to Pi-sufficiency treatments across WT and different transgenic lines for the known Pi-deficiency induced genes of *PHO1* and *SPX* gene family *(SPX1-SPX4)* (Ribot *et al*., 2008; Liu, N *et al*., 2018).

***Supplemental Figure S4. The responsiveness of the bioluminescent sensor plants upon different nutrient deficiency treatments***

(A-D) As indicated in **Fig. 4**, but show the auto-luminescent signals under nitrogen (-N)/phosphate (-P)/potassium (-K) deficiency and without nutrient deficiency (Full) conditions for the other independent lines of 35S (#5), Vector (#7) and Sensor plants (#25).

***Supplemental Figure S5. The responsiveness of the bioluminescent sensor plants upon different phosphate-deficiency treatments***

(A-C) As indicated in **Fig. 5**, but show the Pi-deficiency induced luminescence (A), the mRNA expressions of Phosphate Transporter 1 and luciferase genes (B), and cellular Pi contents (C) for the other independent bioreplicate.

***Supplemental Figure S6. The model performance of different deep-learning methods for image classification***

(A) The prediction performance of various classification models using the image datasets collected from Day11,14 and 18 of bioreplicate#1 and #2 to distinguish the 35S (yellow) and vector (gray) plants (subtask1) as indicated in **Fig. 6A**. (B) As indicated in **Fig. 6B**, but for other indicators of model performances using the datasets collected from Day11,14 and 18 of bioreplicate#1 and #2 to distinguish the plants under Pi sufficient (orange) and deficient conditions (blue) (subtask2). Show the mean and its individual AUROC values from five-fold cross-validation.

***Supplemental Figure S7. The responsiveness of the sensor plants grafted onto chili pepper***

(A,B) As indicated in **Fig. 7**, but shown the luminescence under Pi-deficiency and recovery conditions for the biosensor plants grafted onto the chili pepper plants.

***Supplemental Video 1. Time-lapse luminescent imaging of the sensor plants upon Pi deficiency treatment***

Time-lapse luminescent imaging shows the aerial parts of the transgenic sensor plants grown in vermiculite under Pi sufficient (left) and deficient (right) conditions. (Images collected in the light and darkness were photographed by the DSLR camera every hour from 3pm to 10pm and 11 pm to 7 am (the next day) for ∼two weeks).

***Supplemental Table S1. The test reports of phosphor content of soils.***

***Supplemental Table S2. List of deep-learning methods used in this study.***

***Supplemental Table S3. The list of deep learning models used in this study.***

***Supplemental Table S4. YOLOv9 and SegFormer performance metric*.**

